# Machine-learning classification suggests that many alphaproteobacterial prophages may instead be gene transfer agents

**DOI:** 10.1101/697243

**Authors:** Roman Kogay, Taylor B. Neely, Daniel P. Birnbaum, Camille R. Hankel, Migun Shakya, Olga Zhaxybayeva

## Abstract

Many of the sequenced bacterial and archaeal genomes encode regions of viral provenance. Yet, not all of these regions encode *bona fide* viruses. Gene transfer agents (GTAs) are thought to be former viruses that are now maintained in genomes of some bacteria and archaea and are hypothesized to enable exchange of DNA within bacterial populations. In Alphaproteobacteria, genes homologous to the ‘head-tail’ gene cluster that encodes structural components of the *Rhodobacter capsulatus* GTA (RcGTA) are found in many taxa, even if they are only distantly related to *Rhodobacter capsulatus*. Yet, in most genomes available in GenBank RcGTA-like genes have annotations of typical viral proteins, and therefore are not easily distinguished from their viral homologs without additional analyses. Here, we report a ‘support vector machine’ classifier that quickly and accurately distinguishes RcGTA-like genes from their viral homologs by capturing the differences in the amino acid composition of the encoded proteins. Our open-source classifier is implemented in Python and can be used to scan homologs of the RcGTA genes in newly sequenced genomes. The classifier can also be trained to identify other types of GTAs, or even to detect other elements of viral ancestry. Using the classifier trained on a manually curated set of homologous viruses and GTAs, we detected RcGTA-like ‘head-tail’ gene clusters in 57.5% of the 1,423 examined alphaproteobacterial genomes. We also demonstrated that more than half of the *in silico* prophage predictions are instead likely to be GTAs, suggesting that in many alphaproteobacterial genomes the RcGTA-like elements remain unrecognized.

**Data deposition:** Sequence alignments and phylogenetic trees are available in a **FigShare** repository at DOI 10.6084/m9.figshare.8796419. The Python source code of the described classifier and additional scripts used in the analyses are available via a **GitHub** repository at https://github.com/ecg-lab/GTA-Hunter-v1

## Introduction

Viruses that infect bacteria (phages) are extremely abundant in biosphere (Keen 2015). Some of the phages integrate their genomes into bacterial chromosomes as part of their infection cycle and survival strategy. Such integrated regions, known as prophages, are very commonly observed in sequenced bacterial genomes. For example, Touchon et al. (2016) report that 46% of the examined bacterial genomes contain at least one prophage. Yet, not all of the prophage-like regions represent *bona fide* viral genomes (Koonin and Krupovic 2018). One such exception is a Gene Transfer Agent, or GTA for short (reviewed most recently in Lang et al. [2017] and Grull et al. [2018]). Many of genes that encode GTAs have significant sequence similarity to phage genes, but the produced tailed phage-like particles generally package pieces of the host genome unrelated to the “GTA genome” (Hynes et al. 2012; Tomasch et al. 2018). Moreover, the particles are too small to package complete GTA genome (Lang et al. 2017). Hence, GTAs are different from lysogenic viruses, as they do not use the produced phage-like particles for the purpose of their propagation.

Currently, five genetically unrelated GTAs are known to exist in Bacteria and Archaea (Lang et al. 2017). The best studied GTA is produced by the alphaproteobacterium *Rhodobacter capsulatus* and is referred hereafter as the RcGTA. Since RcGTA’s discovery 45 years ago (Marrs 1974), the genes for the related, or RcGTA-like, elements have been found in many of the alphaproteobacterial genomes (Shakya et al. 2017). For a number of *Rhodobacterales* isolates that carry RcGTA-like genes, there is an experimental evidence of GTA particle production (Fu et al. 2010; Nagao et al. 2015; Tomasch et al. 2018). Seventeen of the genes of the RcGTA “genome” are found clustered in one locus and encode proteins that are involved in DNA packaging and head-tail morphogenesis (Figure 1 and **Supplementary Table S1**). This locus is referred to as a ‘head-tail cluster’. The remaining seven genes of the RcGTA genome are distributed across four loci and are involved in maturation, release and regulation of RcGTA production (Hynes et al. 2016). Since the head-tail cluster resembles a typical phage genome with genes organized in modules similar to those of a λ phage genome (Lang et al. 2017), and since many of its genes have homologs in *bona fide* viruses and conserved phage gene families (Shakya et al. 2017), the cluster is usually designated as a prophage by algorithms designed to detect prophage regions in a genome (Shakya et al. 2017). The RcGTA’s classification as a prophage raises a possibility that some of the ‘*in silico*’-predicted prophages may instead represent genomic regions encoding RcGTA-like elements.

**Figure 1.**
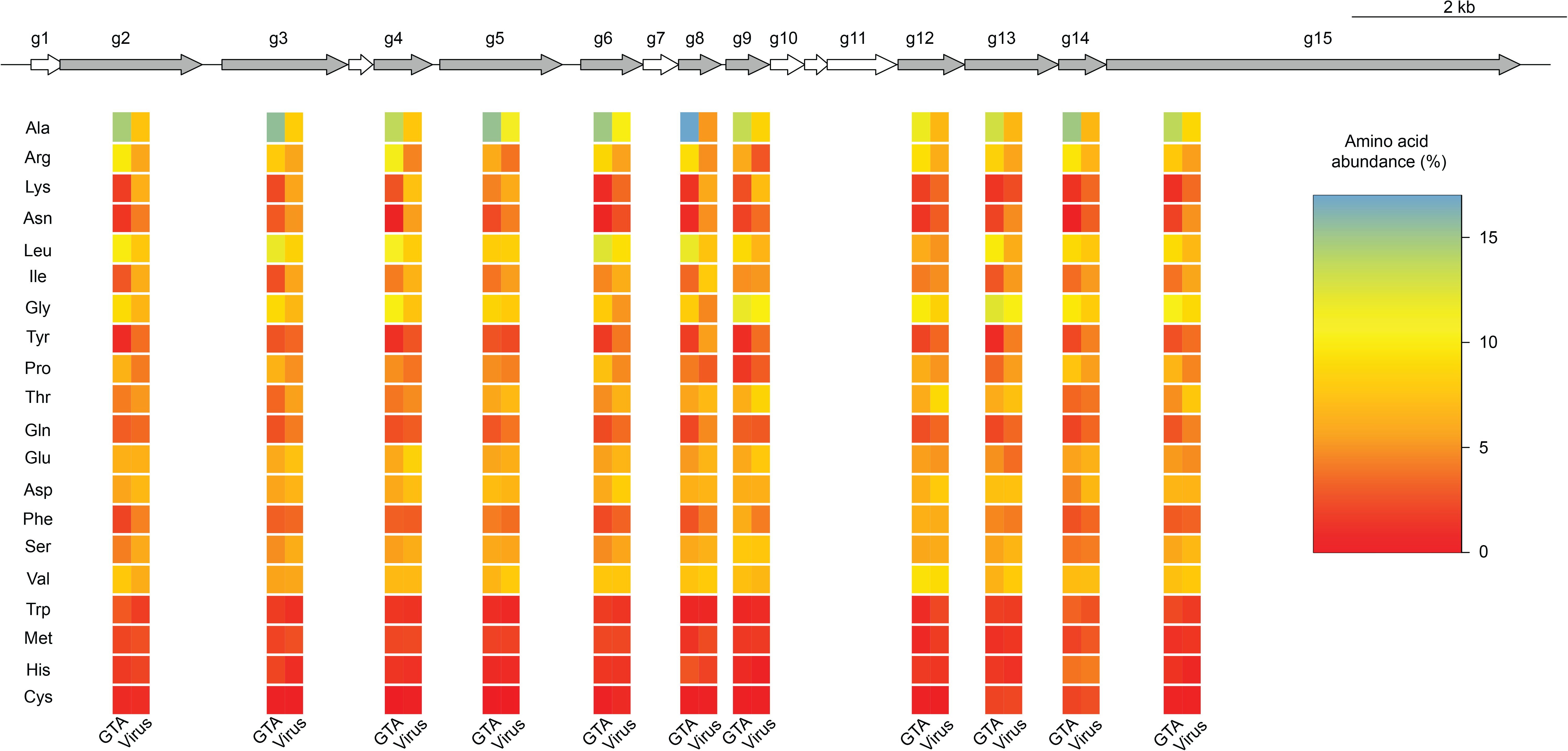
The ‘head-tail’ cluster of the *Rhodobacter capsulatus* GTA “genome” and the amino acid composition of viral and alphaproteobacterial homologs for some of its genes. Genes that are used in the machine learning classification are highlighted in grey. For those genes, the heatmap below a gene shows the relative abundance of each amino acid (rows) averaged across the RcGTA-like and viral homologs that were used in the classifier training (columns). The amino acids are sorted by the absolute difference in the average relative abundance between RcGTA-like and viral homologs, which was additionally averaged across 11 genes. The heatmaps of the amino acid composition in the individual homologs are shown in Supplementary Figure S1.

Presently, to distinguish RcGTA-like genes from the truly viral homologs one needs to examine evolutionary histories of the RcGTA-like and viral homologs and to compare gene content of a putative RcGTA-like element to the RcGTA “genome”. These analyses can be laborious and often require subjective decision making in interpretations of phylogenetic trees. An automated method that could quickly scan thousands of genomes is needed. Notably, the RcGTA-like genes and their viral homologs have different amino acid composition (Figure 1 and Supplementary Figure S1). Due to the purifying selection acting on the RcGTA-like genes at least in the *Rhodobacterales* order (Lang et al. 2012) and of their overall significantly lower substitution rates when compared to viruses (Shakya et al. 2017), we hypothesize that the distinct amino acid composition of the RcGTA-like genes is preserved across large evolutionary distances, and therefore the RcGTA-like genes can be distinguished from their *bona fide* viral homologs by their amino acid composition.

Support vector machine (SVM) is a machine learning algorithm that can quickly and accurately separate data into two classes from the differences in specific features within each class (Cortes and Vapnik 1995). The SVM-based classifications have been successfully used to delineate protein families (e.g., DNA binding proteins (Bhardwaj et al. 2005), G-protein coupled receptors (Karchin et al. 2002), and herbicide resistance proteins (Meher et al. 2019)), to distinguish plastid and eukaryotic host genes (Kaundal et al. 2013), and to predict influenza host from DNA and amino acid oligomers found in the sequences of the flu virus (Xu et al. 2017). During the training step, the SVM constructs a hyperplane that best separates the two classes. During the classification step, data points that fall on one side of the hyperplane are assigned to one class, while those on the other side are assigned to the other class. In our case, the two classes of elements in need of separation are phages and GTAs, while their distinguishing features are several metrics that capture the amino acid composition of the encoding genes.

In this study, we developed, implemented, and cross-validated an SVM classifier that distinguishes RcGTA-like head-tail cluster genes from their phage homologs with high accuracy. We then applied the classifier to 1,423 alphaproteobacterial genomes to examine prevalence of putative RcGTA-like elements in this diverse taxonomic group and to assess how many of the RcGTA-like elements are mistaken for prophages in the *in silico* predictions.

## Materials and Methods

### The Support Vector Machine (SVM) classifier and its implementation

Let’s denote as *u* a homolog of an RcGTA-like gene *g* that needs to be assigned to a class *y*, “GTA” (*y* = − 1) or “virus” (*y* = 1). The assignment is carried out using a weighted soft-margin SVM classifier, which is trained on a dataset of *m* sequences 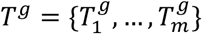 that are homologous to *u* (see **“SVM training data”** section below). The basis of the classification is the *n*-dimensional vector of features ***x*** associated with sequences *u* and 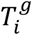 (see **“Generation of sequence features”** section below). Each sequence 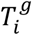 is known to belong to a class *y_i_*.

Using the training dataset *T^g^*, we identify hyperplane that separates two classes as an optimal solution to the objective function:

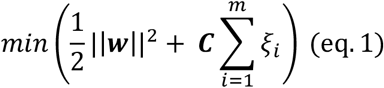

subject to:

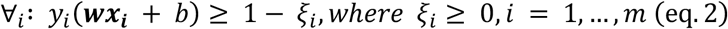

where ***w*** and *b* define the hyperplane *f*(***x***) = ***wx*_*i*_** + *b* that divides the two classes, *ξ_i_* is the slack variable that allows some training data points not to meet the separation requirement, and ***C*** is a regularization parameter, which is represented as an *m* × *m* diagonal matrix. The ***C*** matrix determines how lenient the soft-margin SVM is in allowing for genes to be misclassified: larger values “harden” the margin, while smaller values “soften” the margin by allowing more classification errors. The product ***C****ξ* represents the cost of misclassification. The most suitable values for the ***C*** matrix were determined empirically during cross-validation, as described in the **“Model training, cross validation, and assessment”** section below.

To solve equation 1, we represented this minimization problem in the Lagrangian dual form *L*(*α*):

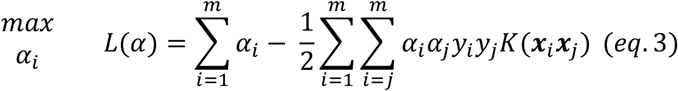

subject to:

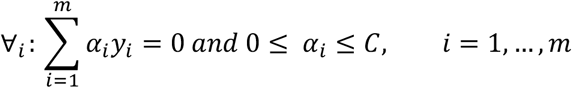

where *K* represents a kernel function. The minimization problem was solved using the convex optimization (CVXOPT) quadratic programming solver (Andersen et al. 2012). The pseudocode of the algorithm for the weighted soft-margin SVM classifier training and prediction is shown in Figure 2.

**Figure 2.**
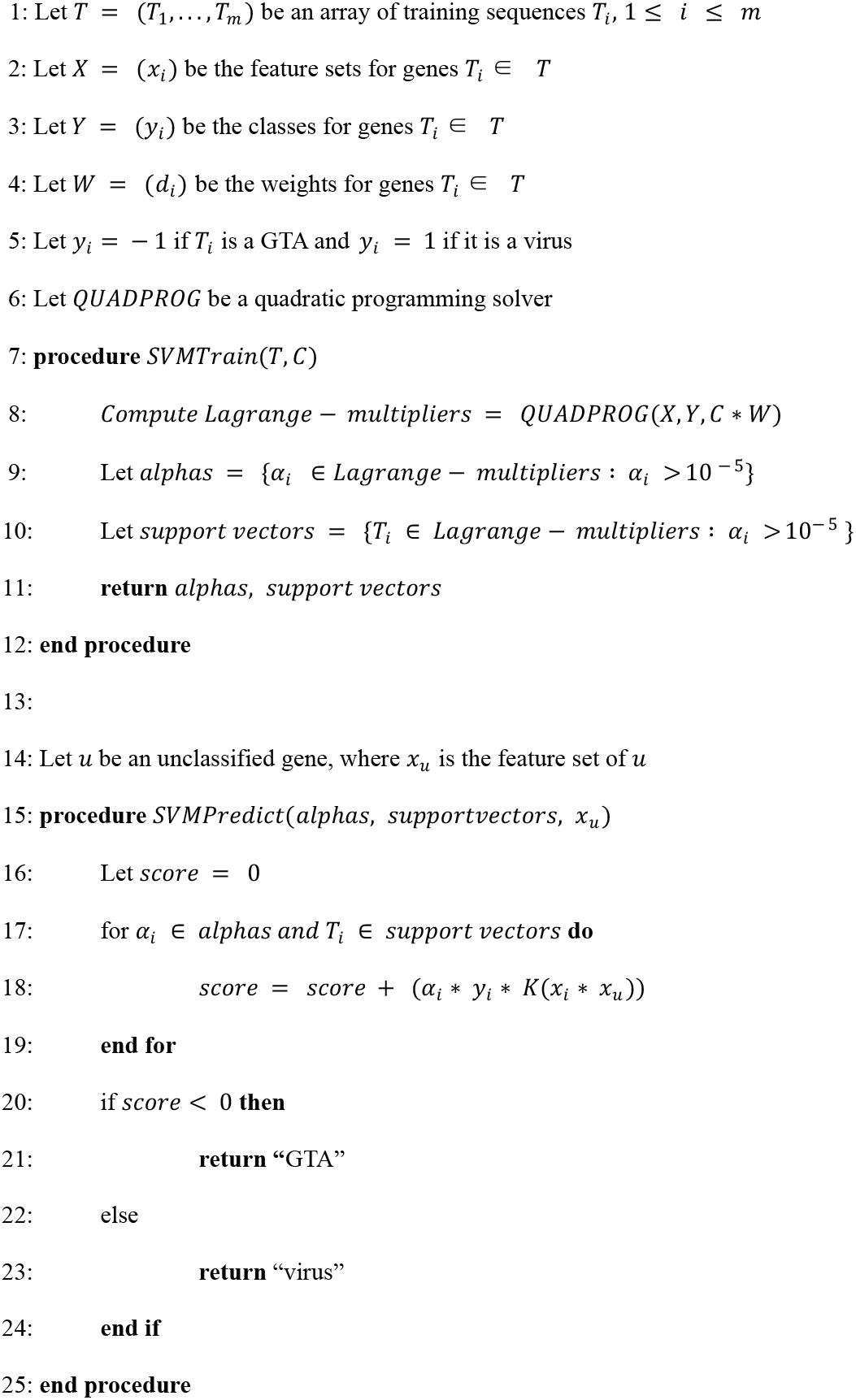
The pseudocode of the SVM classifier algorithm that distinguishes RcGTA-like genes from the ‘true’ viruses. The algorithm is implemented in the GTA-Hunter software package (see **“Software Implementation”** section in **Materials and Methods**).

### SVM training data

To train the classifier, sets of “true viruses” (class *y* = 1) and “true GTAs” (class *y* = −1) were constructed separately for each RcGTA-like gene *g*. To identify the representatives of “true viruses”, amino acid sequences of 17 genes from the RcGTA head-tail cluster were used as queries in BLASTP (E-value < 0.001; query and subject overlap by at least 60% of their length) and PSI-BLASTP searches (E-value < 0.001; query and subject overlap by at least 40% of their length; maximum of six iterations) of the viral RefSeq database release 90 (last accessed in November 2018; accession numbers of the viral entries are provided in **Supplementary Table S2**). BLASTP and PSI-BLAST executables were from the BLAST v. 2.6.0+ package (Altschul et al. 1997). The obtained homologs are listed in **Supplementary Table S3**. Due to few or no viral homologs for some of the queries, the final training sets *T^g^* were limited to 11 out of 17 RcGTA-like head-tail cluster genes (*g2, g3, g4, g5, g6, g8, g9, g12, g13, g14, g15*; see **Supplementary Table S1** for functional annotations of these genes).

As the representatives of the “true GTAs”, we used the RcGTA-like regions that were designated as such via phylogenetic and genome neighborhood analyses by Shakya et al. (2017). To make sure that our “true GTAs” do not contain any other regions, we created a database of the 235 complete alphaprotebacterial genomes that were available in the RefSeq database prior to January 2014 (**Supplementary Table S4**). To identify the representatives of “true GTAs” in this database, amino acid sequences of 17 genes from the RcGTA head-tail cluster (Lang et al., 2017) were used as queries in BLASTP (E-value < 0.001; query and subject overlap by at least 60% of their length) and PSI-BLAST searches (E-value < 0.001; query and subject overlap by at least 40% of their length; maximum of six iterations) of the database. For each genome, the retrieved homologs were designated as an RcGTA-like head-tail cluster if at least 9 of the homologs had no more than 5,000 base pairs between any two adjacent genes. The maximum distance cutoff was based on the observed distances between the neighboring RcGTA head-tail cluster genes. This assignment was determined by clustering of the obtained homologs with the DBSCAN algorithm (Ester et al. 1996) using an in-house Python script (available in a **GitHub** repository; see **“Software Implementation”** section below). The resulting set of 88 “true GTAs” is provided in **Supplementary Table S5** and was verified to represent a subset of RcGTA-like elements that were identified by Shakya et al. (2017).

Since GTA functionality has been extensively studied only in *Rhodobacter capsulatus* SB1003 (Lang et al. 2017) and horizontal gene transfer likely occurred multiple times between the putative GTAs and bacterial viruses (Hynes et al. 2016; Zhan et al. 2016), the bacterial homologs that were both too divergent from other bacterial RcGTA-like homologs and more closely related to the viral homologs were eliminated from the training sets to reduce possible noise in classification. To do so, for each of the 11 trainings sets *T^g^*, all detected viral and bacterial homologs were aligned using MUSCLE v3.8.31 (Edgar 2004) and then pairwise phylogenetic distances were estimated under PROTGAMMAJTT substitution model using RAxML version 8.2.11 (Stamatakis 2014). For each bacterial homolog in a set *T^g^*, the pairwise phylogenetic distances between it and all other bacterial homologs were averaged. This average distance was defined as an outlier distance (*o*) if it satisfied the inequality:

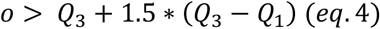

where *Q*_1_ and *Q*_3_ are the first and third quartiles, respectively, of the distribution of the average distances for all bacterial homologs in the training set *T^g^*. If an outlier distance was greater than the shortest distance from it to a viral homolog in the set *T^g^*, the bacterial homolog was removed from the dataset. The alignments, list of removed sequences and the associated calculations are available in the **FigShare** repository.

Additionally, for each gene *g*, the sequences that had the same RefSeq ID (and therefore 100% amino acid identity) were removed from the training data sets. The final number of sequences in each training dataset are listed in Table 1.

**Table 1.** Number of the RcGTA homologs in the “true GTA” and “true virus” training datasets.

| Gene | “true GTAs” | “true viruses” |
| --- | --- | --- |
| *g2* | 69 | 1646 |
| *g3* | 65 | 769 |
| *g4* | 62 | 465 |
| *g5* | 67 | 627 |
| *g6* | 61 | 19 |
| *g8* | 62 | 96 |
| *g9* | 66 | 61 |
| *g12* | 63 | 12 |
| *g13* | 73 | 57 |
| *g14* | 67 | 124 |
| *g15* | 67 | 155 |

### Assignment of weights to the training set sequences

Highly similar training sequences can have an increased influence on the position of the hyperplane, as misclassification of two or more similar sequences can be considered less optimal than misclassification of only one sequence. This could be a problem for our classifier, since there is generally a highly unequal representation of taxonomic groups in the RefSeq database. To correct for this taxonomic bias, a weighting scheme was introduced into the soft-margin of the SVM classifier during training. To do so, sequences in each training set 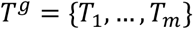 were aligned in MUSCLE v3.8.31 (Edgar 2004) (The alignments are available in the **FigShare** repository). For each pair of sequences in a training set *T^g^*, phylogenetic distances were calculated in RAxML version 8.2.11 (Stamatakis 2014) under the best substitution model (PROTGAMMAAUTO; the selected substitution matrices are listed in the **Supplementary Table S6**). The farthest-neighbor hierarchical clustering method was used to group sequences with distances below a specified threshold *t*. Weight *d_i_* of each sequence in a group was defined as a reciprocal of the number of genes in the group. These weights are used to adjust the cost of misclassification by multiplying *C_ii_* for each sequence *T_i_* by *d_i_*. The most suitable value of *t* was determined empirically during cross-validation, as described in the **“Model training, cross validation, and assessment”** section below.

### Generation of sequence features

To use amino acid sequences in the SVM classifier, each sequence was transformed to an *n*-dimensional vector of compositional features. Three metrics that capture different aspects of sequence composition were implemented: frequencies of “words” of size *k* (*k*-mers), pseudo amino-acid composition (PseAAC), and physicochemical properties of amino acids.

In the first feature type, amino acid sequence of a gene is broken into a set of overlapping subsequences of size *k*, and frequencies of these *n* unique *k*-mers form a feature vector ***x***. Values of *k* equal to 1, 2, 3, 4, 5 and 6 were evaluated for prediction accuracy (see the **“Model training, cross validation, and assessment”** section below).

The second feature type, pseAAC, has n=(20+λ) dimensions and takes into account frequencies of 20 amino acids, as well as correlations of hydrophobicity, hydrophilicity and side-chain mass of amino acids that are λ positions apart in the sequence of the gene (after Chou [2001]), More precisely, PseAAC feature set ***x*** of a sequence of length *L* consisting of amino acids R_1_R_2_…R_L_ is defined as follows:

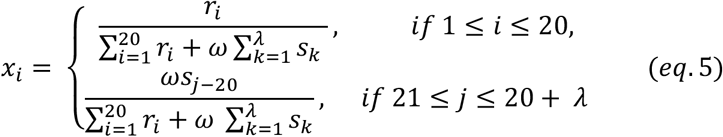

where *r_i_* is the frequency of the *i-*th amino acid (out of 20 possible), *ω* is a weight constant for the order effect that was set to 0.05, and *s_k_* (*k* = 1, …, λ) are sequence order-correlation factors. These factors are defined as

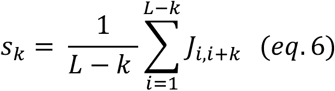

where

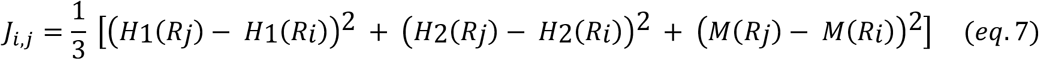

and *H*_1_(*R_i_*), *H*_2_(*R_i_*), and *M*(*R_i_*) denote the hydrophobicity, hydrophilicity, and side-chain mass of amino acid *R_i_*, respectively. The *H*_1_(*R_i_*), *H*_2_(*R_i_*), and *M*(*R_i_*) scores were subjected to a conversion as described in the following equation:

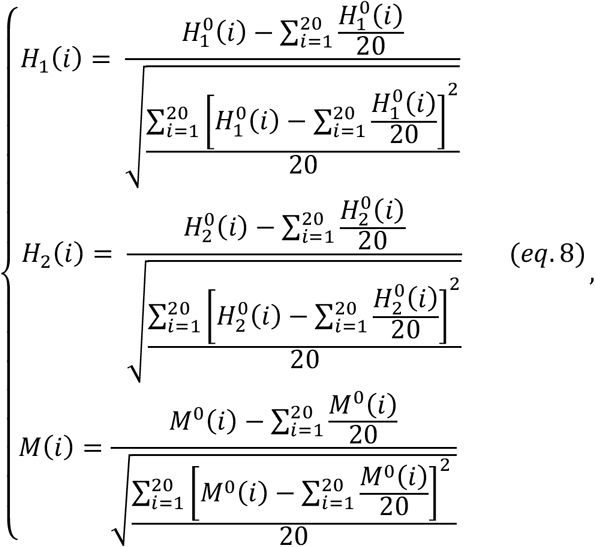

where 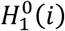 is the original hydrophobicity value of the *i-*th amino acid, 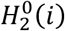 is hydrophilicity value, and 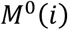 is the mass of its side chain. Values of λ equal to 3 and 6 were evaluated for prediction accuracy (see the **“Model training, cross validation, and assessment”** section below).

The third feature type relies on classification of amino acids into 19 overlapping classes of physicochemical properties (**Supplementary Table S7**; after Kaundal et al. [2013]). For a given sequence, each of its encoded amino acids was counted towards one of the 19 classes, and the overall scores for each class were normalized by the length of the sequence to form *n* = 19-dimensional feature vector ***x***.

### Model training, cross validation, and assessment

For each GTA gene, parameter, and feature type, the accuracy of the classifier was evaluated using a five-fold cross-validation scheme, in which a dataset was randomly divided into five different sub-samples. Four parts were combined to form the training set, while the fifth part was used as the validation set and its SVM-assigned classifications compared to the known classes. This step was repeated five times, so that every set was tested as a known class at least once.

For each class *y* (“GTA” and “Virus”), the results were evaluated by their accuracy scores, defined as the number of correctly classified homologs divided by the total number of homologs that were tested. The cross-validation procedure was repeated ten times to reduce the partitioning bias, and the generated results were averaged, resulting in an Average Accuracy Score (AAS) for each gene and each class. To ensure that “GTA” and “Virus” classes had equal impact on the accuracy assessment, each class was assigned a weight of 0.5. The final, Weighted Accuracy Score (WAS) was calculated as:

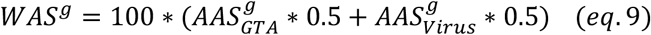

The most suitable “softness” of the SVM margin was determined by trying all possible combinations of several raw diagonal values of the matrix ***C*** (0.01, 0.1, 1, 100, 10000) and the threshold *t* (0, 0.01, 0.02, 0.03, 0.04, 0.05, 0.1). The set of parameters and features that resulted in the highest WAS was defined as the optimal set for a gene *g*. If multiple parameter and feature sets resulted in the equally highest WAS, we applied the following parameter selection criteria, in the priority order listed, until only one parameter set was left: first, we selected parameter set(s) with *k*-mer size that on average performed better than other *k*-mer sizes; second, we avoided parameter set(s) that included PseAAC and physicochemical composition features; third, we selected parameter set(s) with the value of ***C*** that gives the highest average accuracy across the remaining parameter sets; and finally, we opted for the parameter set with the value of *t* that also gives the highest WAS across the remaining parameter sets. Additionally, we evaluated classifier accuracy using the Matthews correlation coefficient (MCC) (Matthews 1975).

### Selection of alphaproteobacterial genomes for testing the presence of RcGTA-like genes

From the alphaproteobacterial genomes deposited to the RefSeq database between January 2014 and January 2019, we selected 636 complete and 789 high-quality draft genomes, with the latter defined as genome assemblies with N50 length >400 kbp. The taxonomy of each genome was assigned using the GTDB-Tk toolkit (Parks et al. 2018). The GTDB assignment is based on the combination of Average Nucleotide Identity (Jain et al. 2018) and phylogenetic placement on the reference tree (as implemented in the *pplacer* program (Matsen et al. 2010)). Three of the 1,425 genomes could not be reliably placed into a known alphaproteobacterial order, and hence were left unclassified. Two of the 1,425 genomes were removed from further analyses due to their classification outside of the Alphaproteobacteria class, resulting in 635 complete and 788 high-quality genomes in our dataset (**Supplementary Table S8**).

### Detection of RcGTA-like genes and head-tail clusters in Alphaproteobacteria

The compiled training datasets of the RcGTA-like genes (see the **“SVM training data”** section) were used as queries in BLASTP (E-value < 0.001; query and subject overlap by at least 60% of their length) searches of amino acid sequences of all annotated genes from the 1,423 alphaproteobacterial genomes. Acquired homologs of unknown affiliation (sequences *u*) were then assigned to either “GTA” or “virus” category by running the SVM classifier with the identified optimal parameters for each gene *g* (Table 2).

**Table 2.** The combinations of features and parameters that showed the highest weighted accuracy score (WAS) in cross-validation. The listed parameter sets were used in predictions of the RcGTA-like genes in 1,423 alphaproteobacterial genomes. See **Materials and Methods** for the procedure on selecting one parameter set in the cases where multiple parameter sets had the identical highest WAS.

| Gene | Weighted Accuracy Score, WAS (%) | Matthews Correlation Coefficient, MCC | k-mer (size) | PseAAC (value of $\lambda$) | Grouping based on physicochemical properties of amino acids | *C* | *t* |
| --- | --- | --- | --- | --- | --- | --- | --- |
| *g2* | 100 | 1 | 2 | -<sup>1</sup> | - | 10000 | 0.02 |
| *g3* | 100 | 1 | 3 | - | - | 10000 | 0.02 |
| *g4* | 100 | 1 | 3 | 3 | - | 10000 | 0.02 |
| *g5* | 100 | 1 | 3 | - | - | 100 | 0.02 |
| *g6* | 95.9 | 0.88 | 4 | - | + | 0.1 | 0.02 |
| *g8* | 99.4 | 0.98 | 2 | 3 | - | 0.1 | 0.03 |
| *g9* | 100 | 1 | 2 | - | - | 100 | 0.1 |
| *g12* | 95.6 | 0.90 | 5 | - | - | 10000 | 0.05 |
| *g13* | 99.1 | 0.98 | 2 | - | - | 100 | 0 |
| *g14* | 99.6 | 0.99 | 6 | 6 | - | 0.01 | 0.03 |
| *g15* | 99.7 | 0.99 | 2 | - | - | 10000 | 0.02 |

<sup>1</sup> throughout the table, “-“ denotes that the feature type was not used

The proximity of the individually predicted RcGTA-like genes in each genome was evaluated by running the DBSCAN algorithm (Ester et al. 1996) implemented in an in-house Python script (available in a **GitHub** repository; see **“Software Implementation”** section below). The retrieved homologs were designated as an RcGTA-like head-tail cluster only if at least 6 of the RcGTA-like genes had no more than 8,000 base pairs between any two adjacent genes. The maximum distance cutoff was increased from the 5,000 base pairs used for the clustering of homologs in the training datasets (see **“SVM Training Data”** section) because the SVM classifier evaluates only 11 of the 17 RcGTA-like head-tail cluster homologs and therefore the distances between some of the identified RcGTA-like genes can be larger.

To reduce the bias arising from the overrepresentation of particular taxa in the estimation of the RcGTA-like cluster abundance in Alphaproteobacteria, the 1,423 genomes were grouped into Operational Taxonomic Units (OTUs) by computing pairwise Average Nucleotide Identity (ANI) using the FastANI v1.1 program (Jain et al. 2018) and defining boundaries between OTUs at the 95% threshold. Since not all OTUs consist uniformly of genomes that were either all with or all without the RcGTA-like clusters, each RcGTA-like cluster in an OTU was assigned a weight of “1/[number of genomes in an OTU]”. The abundance of the RcGTA-like clusters in different alphaproteobacterial orders was corrected by summing up the weighted numbers of RcGTA-like clusters.

### Software Implementation

The above described SVM classifier, generation of sequence features, and preparation and weighting of training data are implemented in a Python program called “GTA-Hunter”. The source code of the program is available via GitHub at https://github.com/ecg-lab/GTA-Hunter-v1. The repository also contains training data for the detection of the RcGTA-like heat-tail cluster genes, examples of how to run the program, and the script for clustering of the detected RcGTA-like genes using the DBSCAN algorithm.

### Assessment of prevalence of the RcGTA-like clusters among putative prophages

Putative prophages in the 1,423 alphaproteobacterial genomes were predicted using the PHASTER web server (Arndt et al. [2016]; accessed in January, 2019). The PHASTER program was chosen due to its solid performance in benchmarking studies (Sousa et al. 2018) and its useful scoring system that ranks predictions based on a prophage region completeness (Song et al. 2019). To restrict our evaluation to likely functional prophages, only predicted prophages with the PHASTER score >90 (i.e., classified as “intact” prophages) were retained for further analyses. The proportion of these predicted intact prophages classified by the GTA-Hunter as “GTA”s was calculated by comparing the overlap between the genomic locations of the predicted intact prophages and the putative RcGTA-like regions.

### Construction of the alphaproteobacterial reference phylogeny

From the set of 120 phylogenetically informative proteins (Parks et al. 2017), 83 protein families that are present in a single copy in >95% of 1,423 alphaproteobacterial genomes were extracted using *hmmsearch* (E-value < 10^−7^) via modified AMPHORA2 scripts (Wu and Scott 2012) (**Supplementary Table S9**). For each protein family, homologs from *Escherichia coli* str. K12 substr. DH10B and *Pseudomonas aeruginosa* PAO1 genomes (also retrieved using *hmmsearch*, as described above) were added to be used as an outgroup in the reconstructed phylogeny. The amino acid sequences of each protein family were aligned using MUSCLE v3.8.31 (Edgar 2004). Individual alignments were concatenated, keeping each alignment as a separate partition in further phylogenetic analyses (Chernomor et al. 2016). The most suitable substitution model for each partition was selected using *ProteinModelSelection.pl* script downloaded from https://github.com/stamatak/standard-RAxML/tree/master/usefulScripts. Gamma distribution with 4 categories was used to account for rate heterogeneity among sites (Yang 1994). The maximum likelihood phylogenetic tree was reconstructed with IQ-TREE v 1.6.7 (Nguyen et al. 2014). One thousand ultrafast bootstrap replicates were used to get support values for each branch (Hoang et al. 2017; Minh et al. 2013). The concatenated sequence alignment in PHYLIP format and the reconstructed phylogenetic tree in Newick format are available in the **FigShare** repository.

### Examination of conditions associated with the decreased fitness of the knock-out mutants of the RcGTA-like head-tail cluster genes

From the three genomes that are known to contain RcGTA-like clusters (*Caulobacter crescentus* NA100, *Dinoroseobacter shibae* DFL-12, and *Phaeobacter inhibens* BS107), fitness experiments data associated with the knock-out mutants of the RcGTA-like head-tail cluster genes were retrieved from the Fitness Browser (Price et al. [2018]; accessed in May, 2019 via http://fit.genomics.lbl.gov/cgi-bin/myFrontPage.cgi). Price et al. (2018) defined gene fitness as the log2 change in abundance of knock-out mutants in that gene during the experiment. For our analyses, the significantly decreased fitness of each mutant was defined as a deviation from the fitness of 0 with a |*t* − *score*| ≥ 4. The conditions associated with the significantly decreased fitness were compared across the RcGTA-like head-tail cluster genes in all three genomes.

## Results

### GTA-Hunter is an effective way to distinguish RcGTA-like genes from their viral homologs

The performance of the developed SVM classifier depends on values of parameters that determine type and composition of sequence features, specify acceptable levels of misclassification, and account for biases in taxonomic representation of the sequences in the training sets. To find the most effective set of parameters, for each of the 11 RcGTA-like head-tail genes with the sufficient number of homologs available (Figure 1; also, see **Materials and Methods** for details) we evaluated the performance of 1,435 different combinations of the parameters using a cross-validation technique (**Supplementary Table S10**).

Generally, the classifiers that only use *k*-mers as the feature have higher median WAS values than the classifiers that solely rely either on physicochemical properties of amino acids or on pseudo amino acid composition (PseAAC) (Supplementary Figure S2 and **Supplementary Table S10**), indicating that the conservation of specific amino acids blocks is important in delineation of RcGTA-like genes from their viral counterparts. However, the WAS values are lower for the large *k*-mer sizes (Supplementary Figure S2), likely due to the feature vectors becoming too sparse. Consequently, parameter combinations with values of *k* above 6 were not tested. The WAS values are also lower for *k=1*, likely due to the low informativeness of the feature. The lowest observed WAS values involve usage of physicochemical properties of proteins as a feature (Supplementary Figure S2 and **Supplementary Table S10**), suggesting the conservation of physicochemical properties of amino acids among proteins of similar function in viruses and RcGTA-like regions despite their differences in the amino acid composition. The more sophisticated re-coding of physicochemical properties of amino acids as the PseAAC feature performs better, but for all genes its performance is worse than the best-performing *k*-mer (Supplementary Figure S2 and **Supplementary Table S10**).

For several genes, the highest value of WAS was obtained with multiple combinations of features and parameter values (**Supplementary Table S10**). Based on the above-described observations of the performance of individual features, we preferred parameter sets that did not include PseAAC and physicochemical composition features, and selected *k*-mer size that on average performed better than other *k*-mer sizes (see **Materials and Methods** for the full description of the parameter selection procedure).

For individual genes, the WAS of the selected parameter set ranges from 95.6 to 100% (Table 2), with 5 out of 11 genes reaching the WAS of 100%. The two genes with the highest WAS below 99% (*g6* and *g12*) have the smallest number of viral homologs available for training (Table 2). Additionally, several viral homologs in the training datasets for *g6* and *g12* genes have smaller phylogenetic distances to “true GTA” homologs than to other “true virus” homologs (**Supplementary Table S11**). As a result, the SVM classifier erroneously categorizes some of the RcGTA-like *g6* and *g12* genes as “viral”, resulting in the reduced classifier efficacy (**Supplementary Table S10**).

Assessment of accuracy using the Matthews correlation coefficient (MCC) generally agrees with the results based on WAS (Table 2 and **Supplementary Table 10**). For 10 out of 11 genes, the set of parameters with the highest WAS also has the highest MCC. For gene *g6*, there are sets of parameters with higher MCC than the MCC for set of parameters with the highest WAS, but the differences among the MCC values are small (**Supplementary Table S10**). Therefore, the combinations of features and parameters chosen using the WAS scheme (Table 2) were selected to classify homologs of the RcGTA genes in the 1,423 alphaproteobacterial genomes (**Supplementary Table S8**).

### GTA-Hunter predicts abundance of RcGTA-like head-tail clusters in Alphaproteobacteria

The 1,423 examined alphaproteobacterial genomes contain 7,717 homologs of the 11 RcGTA genes. The GTA-Hunter classified 6,045 of these homologs as “GTA” genes (**Supplementary Table S12**). However, many genomes are known to contain regions of decaying viruses that may be too divergent to be recognizably “viral” and there is at least one known case of horizontal gene transfer of several GTA genes into a viral genome (Zhan et al. 2016), raising a possibility that some of the predicted “GTA” genes may not be part of “true GTA” genomic regions. To minimize such false positives, we imposed an extra requirement of multiple predicted RcGTA-like genes to be in proximity on a chromosome. Specifically, we called a genomic region the putative RcGTA-like cluster only if it consisted of at least 6 genes classified as “GTA”. We found that the RcGTA-like clusters defined that way are present in one (and only one) copy in 818 of the 1,423 (∼57.5%) examined alphaproteobacterial genomes (**Supplementary Table S13** and Table 3). Uneven taxonomic representation of Alphaproteobacteria among the analyzed genomes may inflate this estimation of the abundance of the GTA-harboring genomes within the class. To correct for this potential bias, 1,423 genomes were grouped into 797 Operational Taxonomic Units (OTUs) based on the average nucleotide identity (ANI) of their genomes (**Supplementary Table S14**). Although indeed some taxonomic groups are overrepresented in the set of 1,423 genomes, in 450 of the 797 OTUs (56.4%) all OTU members contain the putative RcGTA-like clusters (**Supplementary Table S14**).

**Table 3.**
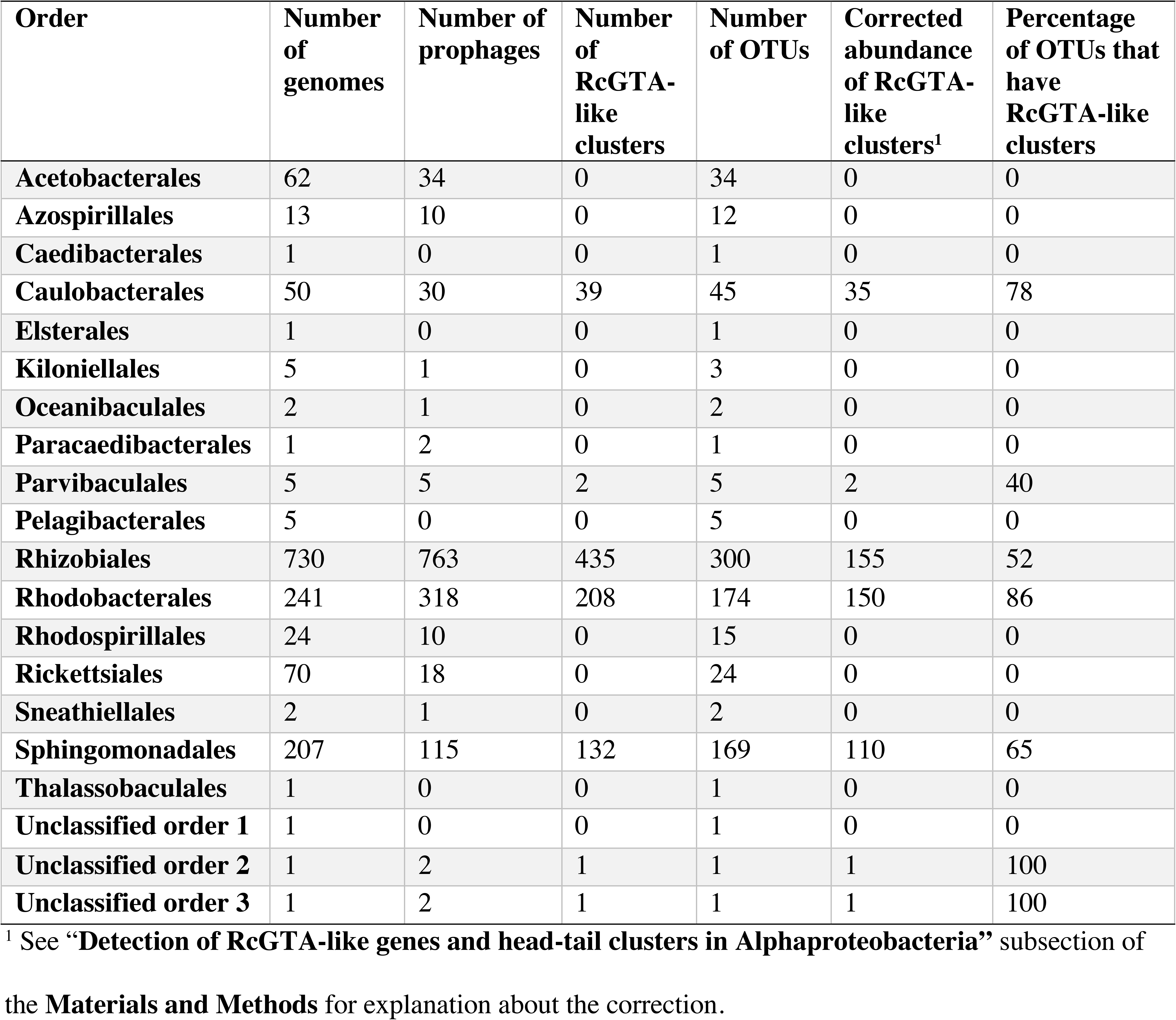
Distribution of prophages and RcGTA-like elements across different orders within class Alphaproteobacteria.

### RcGTA-like clusters are widely distributed within a large sub-clade of Alphaproteobacteria

The 818 genomes with the RcGTA-like gene clusters detected in this study are not evenly distributed across the class (Table 3), but are found only in a clade that includes seven orders (clade 4 in Figure 3). Overall, 66% of the examined OTUs within the clade 4 are predicted to have an RcGTA-like cluster (Table 3). RcGTA-like clusters are most abundant in clade 6 (Figure 3), a group that consists of the orders *Rhodobacterales* and *Caulobacterales* (Table 3).

**Figure 3.**
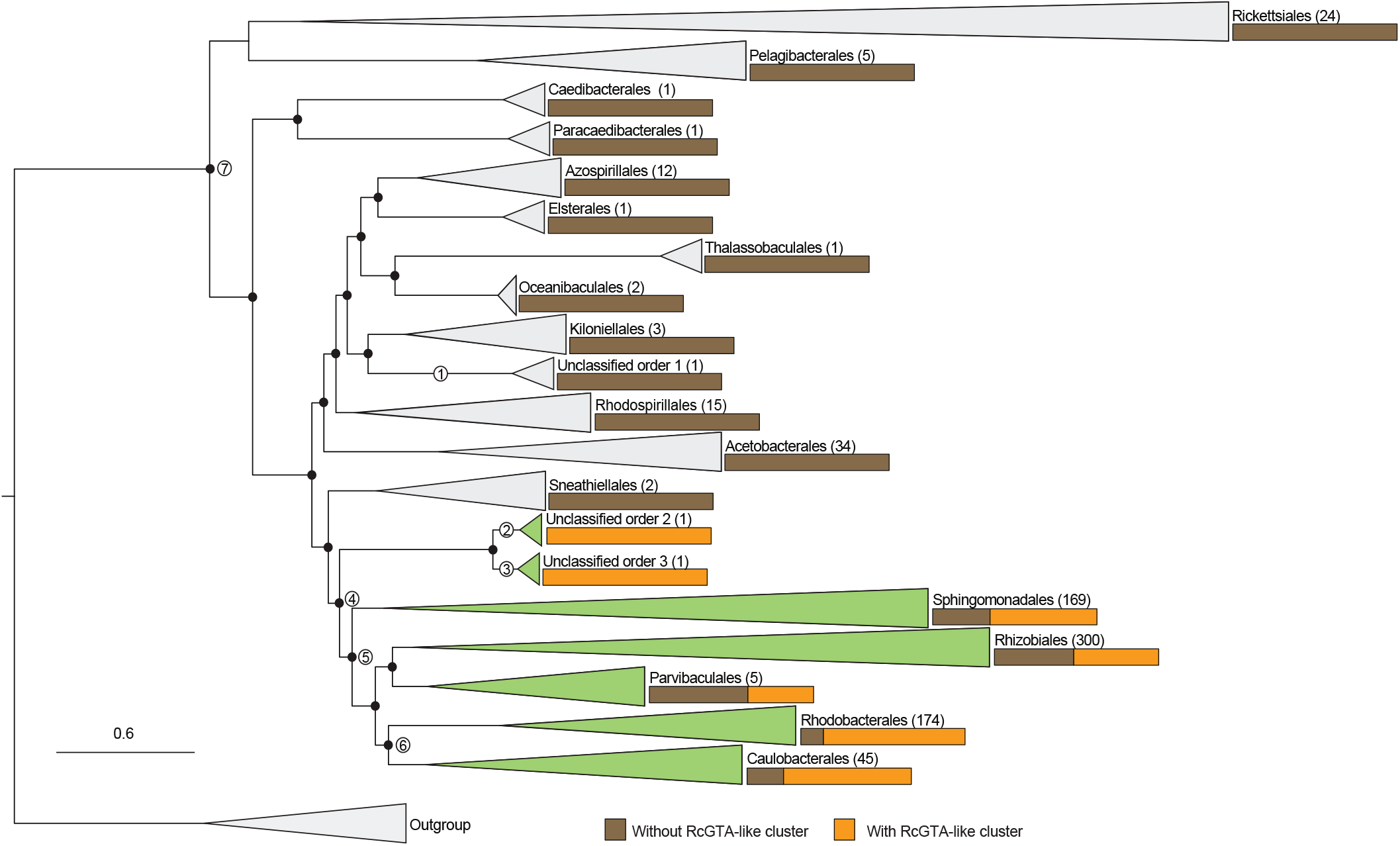
Distribution of the detected RcGTA-like clusters across the class Alphaproteobacteria. The presence of RcGTA-like clusters is mapped to a reference phylogenetic tree that was reconstructed from a concatenated alignment of 83 marker genes (See **Materials and Methods** and **Supplementary Table S9**). The branches of the reference tree are collapsed at the taxonomic rank of “order”, and the number of OTUs within the collapsed clade is shown in parentheses next to the order name. Orange and brown bars depict the proportion of OTUs with and without the predicted RcGTA-like clusters, respectively. The orders that contain at least one OTU with an RcGTA-like cluster are colored in green. Nodes 1, 2 and 3 mark the last common ancestors of the unclassified orders. Node 4 marks the lineage where, based on this study, the RcGTA-like element should have already been present. Nodes 5 and 7 mark the lineages that were previously inferred to represent last common ancestor of the RcGTA-like element by Shakya et al. (2017) and Lang and Beatty (2007), respectively. Node 6 marks the clade where RcGTA-like elements are the most abundant. The tree is rooted using homologs from *Escherichia coli* str. K12 substr. DH10B and *Pseudomonas aeruginosa* PAO1 genomes. Branches with ultrafast bootstrap values >= 95% are marked with black circles. The scale bar shows the number of substitutions per site. The full reference tree is provided in the **FigShare** repository.

Although the two unclassified orders that contain RcGTA-like clusters are represented by only two genomes (clades 2 and 3 in Figure 3), their position on the phylogenetic tree of Alphaproteobacteria suggests that the RcGTA-like element may have originated earlier than was proposed by Shakya et al. (2017) (clade 5 on Figure 3). Given that RcGTA-like head-tail cluster genes are readily detectable in viral genomes, it is unlikely that the RcGTA-like clusters remained completely undetectable in the examined genomes outside of the clade 4 due to the sequence divergence. Therefore, an RcGTA-like element was unlikely to be present in the last common ancestor of all Alphaproteobacteria (clade 7 on Figure 3), which was suggested when only a limited number of genomic data was available (Lang and Beatty 2007).

### Most of the detected RcGTA-like clusters can be mistaken for prophages

Among the 818 detected RcGTA-like clusters, the functional annotations of the 11 examined genes were similar to the prophages and none of them refer to a “gene transfer agent” (data not shown). Since at least 11 of the 17 RcGTA head-tail cluster genes have detectable sequence similarity to viral genes (**Supplementary Table S3**), it is likely that, if not recognized as GTAs, many of the putative RcGTA-like clusters will be designated as “prophages” in genome-wide searches of prophage-like regions. To evaluate this hypothesis, we predicted prophages in the set of 1,423 alphaproteobacterial genomes, and limited our analyses to the predicted prophage regions that are more likely to be functional integrated viruses (‘intact’ prophages; see **Materials and Methods** for the criteria). Indeed, of the 1,235 ‘intact’ prophage regions predicted in the clade 4 genomes, 664 (54%) coincide with the RcGTA-like clusters (Figure 4). Conversely, 664 out of 818 of the predicted RcGTA-like clusters (81%) are classified as intact prophages. Of the 351 RcGTA-like clusters that contain *all* 11 examined genes, 323 (92%) are classified as intact prophages.

**Figure 4.**
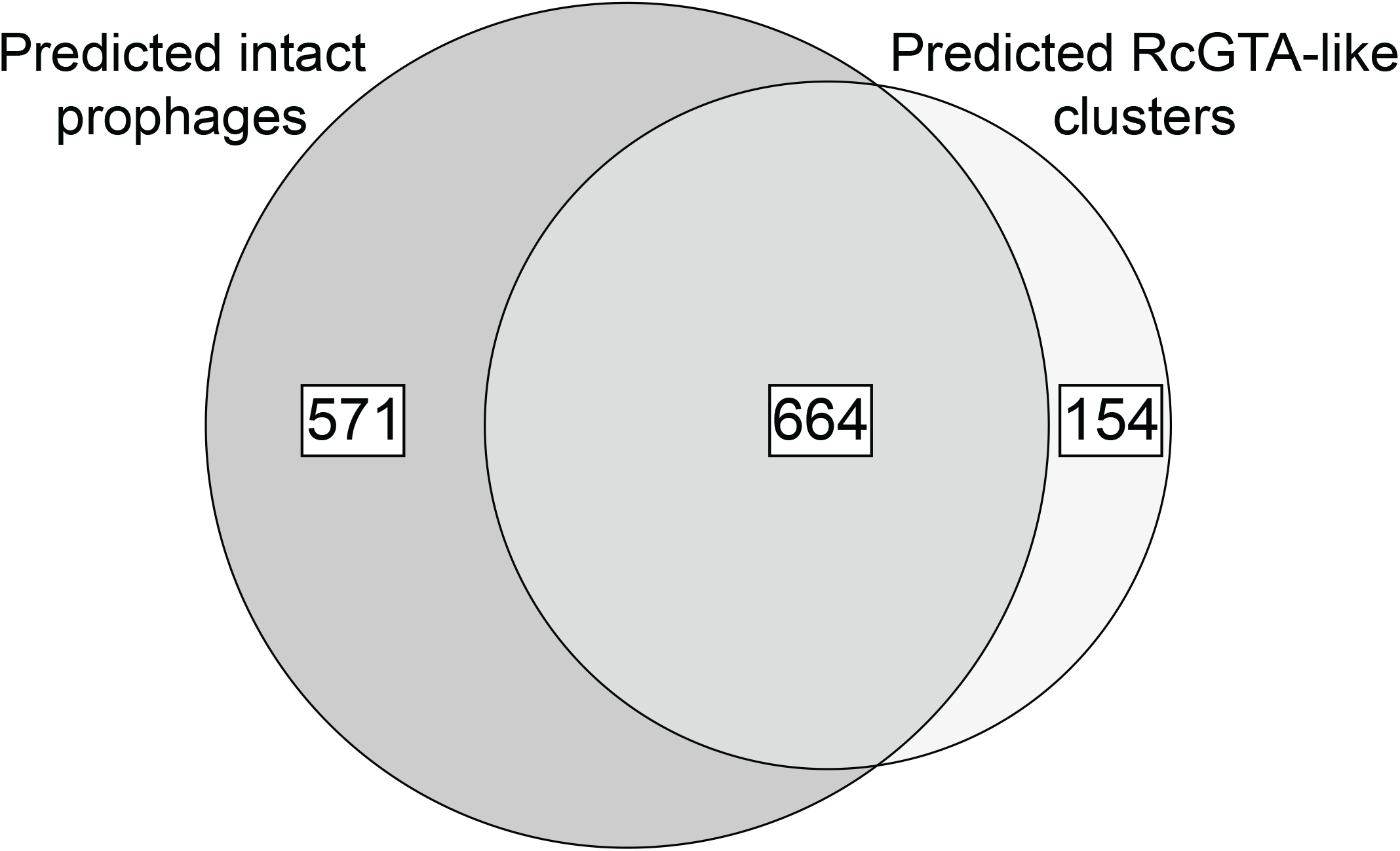
An overlap between prophage and GTA predictions. The “predicted RcGTA-like clusters” set refers to the GTA-Hunter predictions, while the “predicted intact prophages” set denotes predictions made by the PHASTER program (Arndt et al. 2016) on the subset of the genomes that are found within clade 4 (Figure 3).

Interestingly, within 818 genomes that contain RcGTA-like clusters, the average number of predicted intact prophages is 1.23 per genome (Figure 5), which is significantly higher than 0.51 prophages per genome in genomes not predicted to contain RcGTA-like clusters (p-value < 0.22 * 10^−17^; Mann-Whitney U test). If the 664 RcGTA-like regions classified as intact prophages are removed from the genomes that contain them, the average number of predicted ‘intact’ prophages per genome drops to 0.42 (Figure 5) and the difference becomes insignificant (p-value = 0.1492; Mann-Whitney U test). This analysis suggests that an elevated number of the observed predicted prophage-like regions in some alphaproteobacterial genomes may be due to the presence of unrecognized RcGTA-like elements.

**Figure 5.**
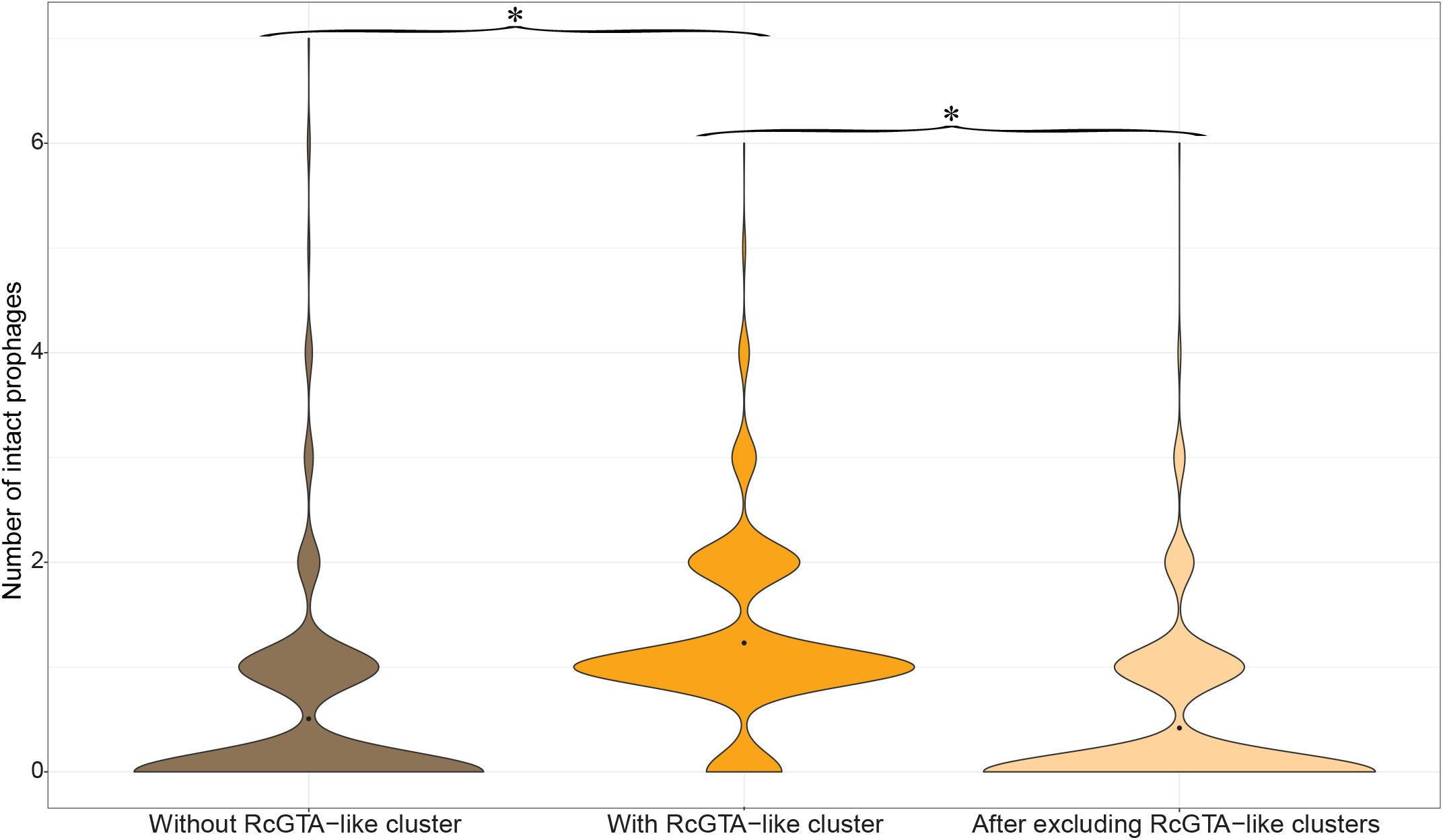
The number of predicted ‘intact’ prophages in alphaproteobacterial genomes. The 1,423 genomes were divided into two groups: those without GTA-Hunter-predicted RcGTA-like clusters (in brown) and those with these RcGTA-like clusters (in dark orange). For the latter group, the number of prophages was re-calculated after the RcGTA-like clusters that were designated as prophages were removed (in light orange). The distribution of the number of predicted intact prophages within each dataset is shown as a violin plot with the black point denoting the average value. The datasets with significantly different average values are denoted by asterisks (p < 0.001; Mann-Whitney U test).

## Discussion

Our study demonstrates that RcGTA-like and *bona fide* viral homologs can be clearly separated from each other using a machine learning approach. The highest accuracy of the classifier is achieved when it primarily relies on short amino acid *k*-mers present in the examined genes. This suggests that the distinct primary amino acid composition of the RcGTA-like and truly viral proteins is what allows the separation of the two classes of elements (Figure 1). However, the cause of the amino acid preferences of the RcGTA-like genes, and especially enrichment of the encoded proteins in alanine and glycine amino acids (Figure 1), remains unknown. Given the structure of the genetic code, the skewed amino acid composition may be the driving force behind the earlier described significantly higher %G+C of the genomic region encoding the RcGTA-like head-tail cluster than the average %G+C in the host genome (Shakya et al. 2017). Regardless of the cause of the skewed amino acid composition, the successful identification of the putative RcGTA-like elements in alphaproteobacterial taxa only distantly related to *Rhodobacter capsulatus* (clade 4 in Figure 3) suggests that the selection to maintain these elements likely extends beyond the *Rhodobacterales* order. Nevertheless, whether these putative elements indeed encode GTAs, as we currently understand them, remains to be experimentally validated.

The benefits associated with the GTA production that would underlie the selection to maintain them remain unknown. In a recently published high-throughput screen for phenotypes associated with specific genes (Price et al. 2018), knockout of the RcGTA-like genes in the three genomes that encode the RcGTA-like elements resulted in decreased fitness of the mutants (in comparison to the wild type) under some of the tested conditions (**Supplementary Table S15**). Interestingly, the conditions associated with the most statistically significant decreases in fitness correspond to the growth on non-glucose sugars, such as D-Raffinose, β-Lactose, D-Xylose and m-Inositol. Overall, carbon source utilization is the most common condition that elicits statistically significant fitness decreases in the mutants. The RcGTA production was also experimentally demonstrated to be stimulated by carbon depletion (Westbye et al. 2017). Further experimental work is needed to identify the link between the RcGTA-like genes expression and carbon utilization. Conversely, absence of the RcGTA-like elements in some of the clade 4 genomes (Figure 3) indicates that in some ecological settings RcGTA-like elements are either deleterious or “useless” and thus their genes were either purged from the host genomes (if RcGTA-like element evolution is dominated by vertical inheritance) or not acquired (if horizontal gene transfer plays a role in the RcGTA-like element dissemination).

Previous analyses inferred that RcGTA-like elements had evolved primarily vertically, with few horizontal gene exchanges between closely related taxa (Hynes et al. 2016; Lang and Beatty 2007; Shakya et al. 2017). Under this hypothesis, the distribution of the RcGTA-like head-tail clusters in alphaproteobacterial genomes suggests that RcGTA-like element originated prior to the last common ancestor of the taxa in clade 4 (Figure 3). This places the origin of the RcGTA-like element to even earlier timepoint than the one proposed in Shakya et al. (2017). However, it should be noted that our inference is sensitive to the correctness of the inferred relationships of taxa within the alphaproteobacterial class, which remain to be disputed due to compositional biases and unequal rates of evolution of some alphaproteobacterial lineages (Munoz-Gomez et al. 2019). The most recent phylogenetic inference that takes into account these heterogeneities (Munoz-Gomez et al. 2019) is different from the reference phylogeny shown in Figure 3. Relevant to the evolution of RcGTA-like elements, on the phylogeny in Munoz-Gomez et al. (2019) the order Pelagibacterales is located within the clade 4 instead of being one of the early-branching alphaproteobacterial orders (Figure 3). No RcGTA-like clusters were detected in Pelagibacterales, although in our analyses the order is represented by only five genomes. Better sampling of genomes within this order would be needed either to show a loss of the RcGTA-like element in this order or to re-assess the hypothesis about origin and transmission of the RcGTA-like elements within Alphaproteobacteria.

Genes in the detected RcGTA-like head-tail clusters remain mainly unannotated as “gene transfer agents” in GenBank records, and therefore they can be easily confused with prophages. For example, recently described “conserved prophage” in *Sphingomonadales* (Viswanathan et al. 2017) is predicted to be an RcGTA-like element by GTA-Hunter. Incorporation of a GTA-Hunter-like machine learning classification into an automated genome annotation pipeline will help improve quality of the gene annotations in GenBank records and facilitate discovery of GTA-like elements in other taxa. Moreover, application of the presented GTA-Hunter program is not limited to the detection of the RcGTA-like elements. With appropriate training datasets, the program can be applied to the detection of GTAs that do not share evolutionary history with the RcGTA (Lang et al. 2017) and of other elements that are homologous to viruses or viral sub-structures, such as type VI secretion system (Leiman et al. 2009) and encapsulins (Giessen and Silver 2017).

## Acknowledgements

This work was supported by the National Science Foundation [NSF-DEB 1551674 to O.Z.]; the Simons Foundation Investigator in Mathematical Modeling of Living Systems [327936 to O.Z.]; Dartmouth Dean of Faculty start-up funds to O.Z.; and Dartmouth James O. Freedman Presidential Scholarship to T.N.

## Supplementary Figure Legends and Table Captions

**Supplementary Figure S1.**
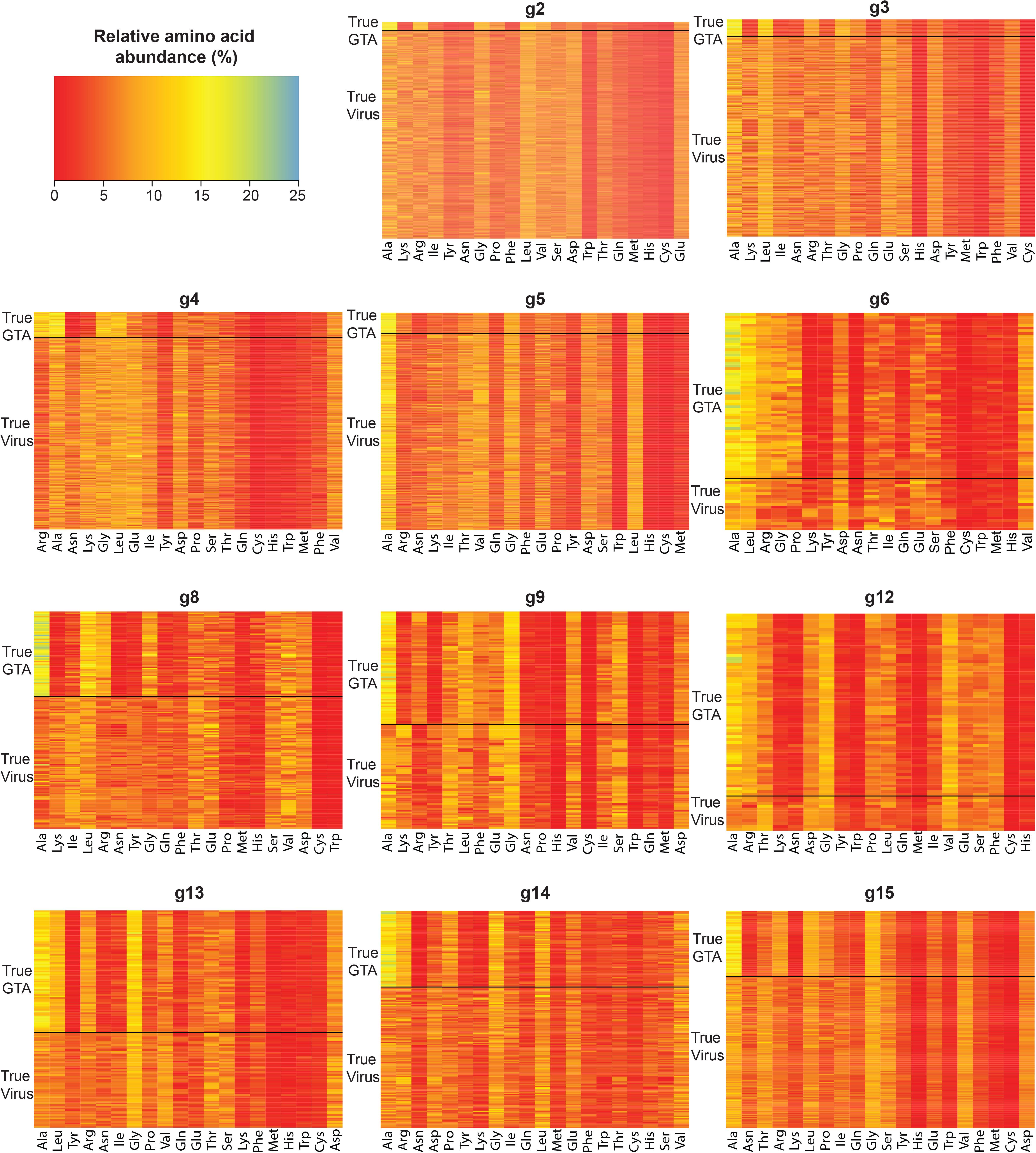
The amino acid composition of viral and alphaproteobacterial homologs of the 11 RcGTA genes. These homologs were used in the training and cross-validation of the SVM classifier. Each heatmap corresponds to one of the 11 genes (see **Supplementary Table S1** for the functional annotations of the genes). Each row in a heatmap corresponds to an individual homolog of the RcGTA gene. The homologs from viruses and alphaproteobacterial are separated by the black line and labeled as “True Virus” and “True GTA”, respectively. The heatmap shows the relative abundance of each amino acid within a homolog. For each gene, the amino acids on the X axis are sorted by the absolute difference in the average relative abundance between “true viruses” and “true GTAs” (from highest on the left to the lowest on the right).

**Supplementary Figure S2.**
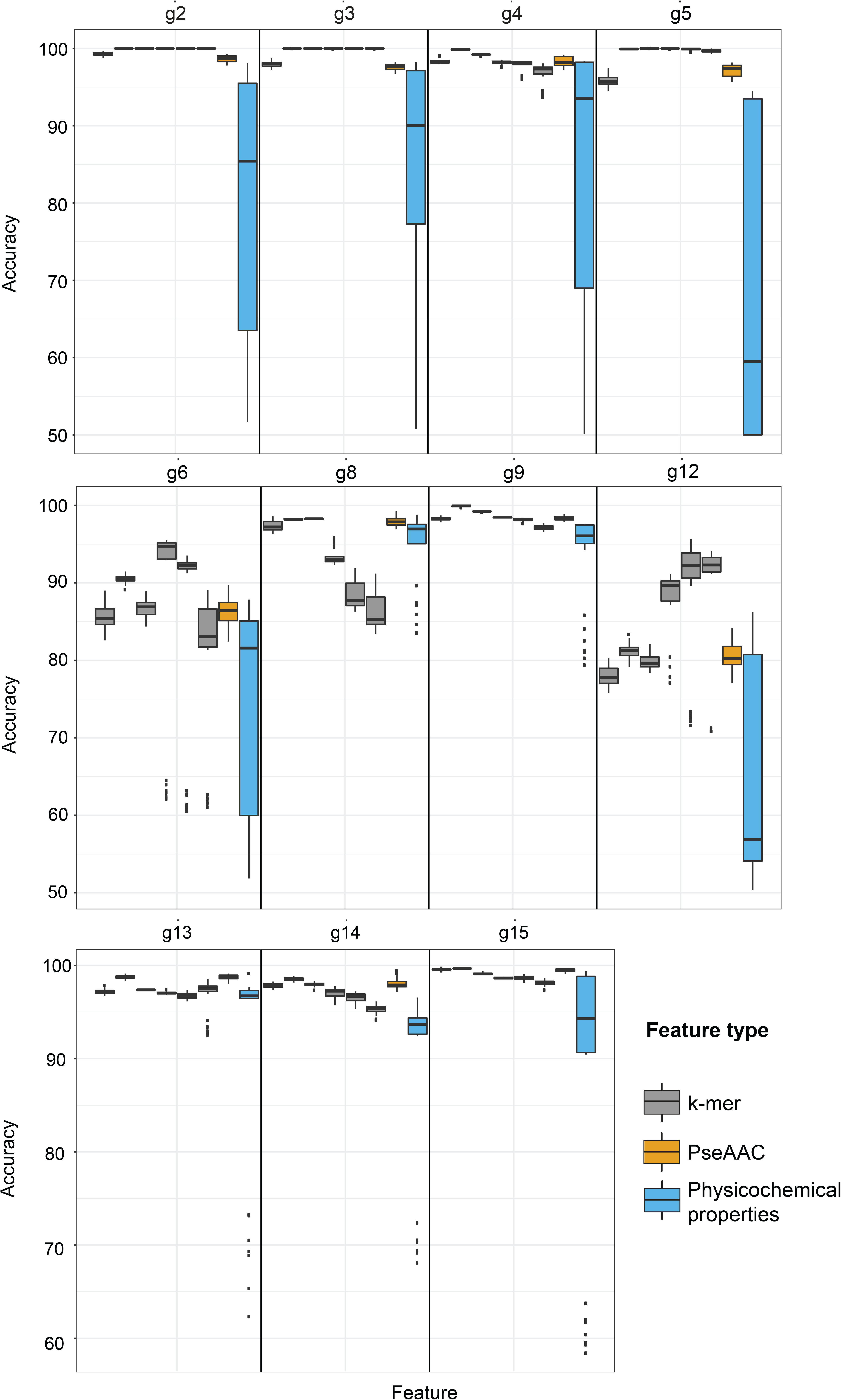
The weighted accuracy scores for different types of features. The boxplots for the three feature types are color coded. The data for six examined *k*-mer sizes (1, 2, 3, 4, 5, 6) are shown from the left to the right on the graphs. Each boxplot shows a median value bounded by the first and third quartiles, and the whiskers depict a deviation that was calculated using the 1.5*InterQuartile Range rule. Outliers are shown as dots.

**Supplementary Table S1. Functional annotations of the ‘head-tail’ cluster genes of the *Rhodobacter capsulatus* gene transfer agent.**

**Supplementary Table S2. List of the 7,995 viral assemblies used to find RcGTA homologs for the training datasets.**

**Supplementary Table S3. List of 1,939 viruses with at least one detected RcGTA homolog.** The data in the columns show the accession numbers of these homologs.

**Supplementary Table S4. List of 235 alphaproteobacterial genomes used to find RcGTA-like clusters for the training datasets.**

**Supplementary Table S5. List of 88 alphaproteobacterial RcGTA-like clusters detected in 85 genomes.** The data in the columns show the accession numbers of these homologs.

**Supplementary Table S6. Substitution matrices that were used to generate pairwise phylogenetic distances within training datasets.**

**Supplementary Table S7. Grouping of amino acids into classes based on their physicochemical properties (after Kaundal et al., 2013).**

**Supplementary Table S8. List of 1,423 alphaproteobacterial genomes used for testing the presence of RcGTA-like genes and regions.**

**Supplementary Table S9. Information about 83 marker genes that were used to reconstruct reference phylogeny of Alphaproteobacteria.**

**Supplementary Table S10. Summary of the classifier cross-validation.** Results for each gene are shown in separate tabs. Each row represents one of the 1,435 tested combinations of the parameters (columns A-E), number and fraction of correctly classified homologs averaged across 10 replicates (columns F-I), and the overall weighted accuracy score (column J) and the Matthews Correlation Coefficient (Column K) of the parameter combination. When a feature was not used, the value of the parameter shown in columns A-C is set to 0.

**Supplementary Table S11. Phylogenetic distances of the “truly viral” homologs of the genes *g6* and *g12* to “true GTAs” and to other “true viruses” in the training datasets**. Data for the *g6* and *g12* homologs are shown in separate tabs. Viral homologs that are more closely related to “true GTAs” than to other “true viruses” are highlighted in yellow.

**Supplementary Table S12. Summary of the alphaproteobacterial RcGTA homologs’ classification.**

**Supplementary Table S13. Information about the 818 detected RcGTA-like clusters.** Data in columns D-N correspond to the RefSeq accession numbers of the encoded proteins.

**Supplementary Table S14. Presence of the RcGTA-like clusters in the reconstructed alphaproteobacterial Operational Taxonomic Units (OTUs).**

**Supplementary Table S15. Results of the fitness experiments with the knock-out mutants of the RcGTA-like head-tail cluster genes in three alphaproteobacterial genomes.** The data was retrieved from the Fitness Browser (Price et al 2018). Each row corresponds to a separate experiment, in which the specified gene was knocked out (column B) and decreased fitness (columns E and F) was associated with a specific condition (column D). The conditions are classified into groups (column C). Rows corresponding to the “carbon source” group are highlighted in yellow. This group is the most common among the listed experiments and is found in experiments associated with each of the three genomes. For description of conditions, refer to the Fitness Browser (Price et al 2018).

